# Ultra-fast Identity by Descent Detection in Biobank-Scale Cohorts using Positional Burrows–Wheeler Transform

**DOI:** 10.1101/103325

**Authors:** Ardalan Naseri, Xiaoming Liu, Shaojie Zhang, Degui Zhi

## Abstract

With the availability of genotyping data of very large samples, there is an increasing need for tools that can efficiently identify genetic relationships among all individuals in the sample. One fundamental measure of genetic relationship of a pair of individuals is *identity by descent* (IBD), chromosomal segments that are shared among two individuals due to common ancestry. However, the efficient identification of IBD segments among a large number of genotyped individuals is a challenging computational problem. Most existing methods are not feasible for even thousands of individuals because they are based on pairwise comparisons of all individuals and thus scale up quadratically with sample size. Some methods, such as GERMLINE, use fast dictionary lookup of short seed sequence matches to achieve a near-linear time efficiency. However, the number of short seed matches often scales up super-linearly in real population data.

In this paper we describe a new approach for IBD detection. We take advantage of an efficient population genotype index, Positional BWT (PBWT), by Richard Durbin. PBWT achieves linear time query of perfectly identical subsequences among all samples. However, the original PBWT is not tolerant to genotyping errors which often interrupt long IBD segments into short fragments. We introduce a randomized strategy by running PBWTs over random projections of the original sequences. To boost the detection power we run PBWT multiple times and merge the identified IBD segments through interval tree algorithms. Given a target IBD segment length, RaPID adjust parameters to optimize detection power and accuracy.

Simulation results proved that our tool (RaPID) achieves almost linear scaling up to sample size and is orders of magnitude faster than GERMLINE. At the same time, RaPID maintains a detection power and accuracy comparable to existing mainstream algorithms, GERMLINE and IBDseq. Running multiple times with various target detection lengths over the 1000 Genomes Project data, RaPID can detect population events at different time scales. With our tool, it is feasible to identify IBDs among hundreds of thousands to millions of individuals, a sample size that will become reality in a few years.

## 1. Introduction

With the recent advances in technology, enormous amounts of population genotype data are being generated from genome-wide SNP array or whole genome sequencing. The 1000 Genome Project [1] has already sequenced more than 2500 individuals. UK biobank [2] will soon release genotype data of around 500,000 individuals. We expect databases from hundreds of thousands up to millions of genotyped data will be abundantly available in the near future. However, the increasing volume of genetic data does not automatically translate into better insight into precision medicine and population genetics. Current methods cannot efficiently detect genetic relationships among all individuals, therefore advanced methods are needed.

A fundamental measure of the genetic relationship is *identity by descent* (IBD). IBD is defined as chromosomal segments shared among two individual chromosomes which have been passed down by a common ancestor. Diploid organisms have two sets of chromosomes and the sets are recombined and passed down to the offspring. As a result, the length of an IBD segment decreases after each generation. The length of an IBD segment after n generations is expected to be 1/2n Morgans because of 2n meiosis [3]. For example after 20 generations the expected IBD length between two related individuals is 1/40 Morgans or 2.5 centimorgans (cM). The length of an IBD due to an ancient ancestor is expected to be even shorter. Short IBDs can be difficult to detect as two random short chromosome segments can have identical sequences – often called *identical by state* (IBS) – by chance. The IBD segments may also not be exactly the same due to the mutation events cumulated over time.

Identification of IBDs is of great interest and has many applications in genetics [4]. Considering the correlations between individuals which can be computed using IBD segments, it is essential to prevent false-positive signals in genome-wise association studies [5]. The IBD segments can be used to determine the pedigree relationship. Pedigree-based relationships can help us to correct the variance of association studies [6]. The detected IBDs can also be directly used in a technique called IBD mapping which detects signals of disease-causing variants in population samples. IBD mapping tests whether cases share more segments of IBDs around a possible variant than controls [7]. Analysis of IBDs in pedigree-based relationships is also important for linkage mapping [8]. The rate at which genes separate and recombine can be used to map the distance between genes. Linkage mapping is crucial for identification of the location of genes that cause genetic diseases. Recent IBDs can also be used for genotype-imputation and haplotype-phase inference [9]. Detection of ancient IBDs is useful in investigating the population structure [10, 11]. By looking at the number of shared IBD segments, we can learn about the overlapping genealogies and the number of common ancestors from which two individuals share genetic sequences [12]. IBD segments can also be used to compute genetic similarity and infer migration patterns [13].

Several methods have been proposed for IBD detection. The proposed methods mostly rely on pairwise comparison of all individuals. One of the classical IBD detection tools is PLINK [14]. PLINK computes for each pair of individuals an IBS score and uses a hidden Markov Model (HMM) method to find IBDs. The hidden IBD state is estimated by the computed IBS sharing and genome-wide level of relatedness. PLINK is slow because even the first step of computing all IBS scores requires *O*(*n*^2^) operations for *n* samples. Beagle IBD [3] computes the ratio of the probability of IBD based on a genotyping error model and the probability of IBS based on haplotype frequencies. Beagle IBD is still computationally expensive and cannot be applied to all pairs of SNPs data without extensive computational resources. FastIBD [15] accounts for haplotype frequencies and uncertain haplotype phase. Although FastIBD is reported to be 1000 times faster than Beagle IBD method on a single core of an Intel Xeon E5620 running at 2.4 GHz, its running time is still a bottleneck for very large populations. PARENTE [16] first divides the entire sequence into fixed sized windows and then computes the likelihood ratio for the entire window. This approach provides faster IBD detection (~10 times faster than FastIBD) but it is still not fast enough for large populations. The new version of PARENTE (PARENTE2) [17] also employs a window-based approach, but the windows contain non-consecutive, and randomly selected markers. PARENTE2 also aggregates multiple haplotypic models to estimate the likelihood in order to increase accuracy. Using PARENTE2 with a filter called SpeeDB [18] resulted in ~10 pairs/second. Consider a population with ~10,000 individual, PARENTE2 will require ~50 million comparisons or 58 days. IBDseq [19] uses a probabilistic model based on allele frequencies and expected error rate accounting for genotyping errors but its running time is a major bottleneck when applying this algorithm for large populations. In general, pairwise comparison of all individuals and their allele sites is not scalable for hundreds of thousands of individuals and millions of variant sites.

One of the fastest IBD detection methods is GERMLINE [20]. GERMLINE avoids pairwise comparison by adopting a seed-and-extension approach. It first builds a dictionary of short subsequences (words) in all individuals, and then find all exact matches of words, *seeds*, by hash table lookup. The detected seed matches are then extended and merged allowing mismatches. The overall assumption of GERMLINE is that the number of seed matches between an individual and others is constant and thus the running time of GERMLINE grows linearly with the number of samples. However, population genotype sequence is not a random sequence. Some short sequences (haplotypes) may persist in a large number of individuals and thus GERMLINE may deviate from its idealized linear behavior.

An analogy to IBD detection can be drawn with the DNA sequence search (see Table 1). In order to search for DNA sequences in a large reference genome, instead of performing the time-consuming sequence alignment (Smith-Watermann algorithm [21]), keyword-based search approaches like BLAST [22] have proved to be more efficient. With the increasing number of queries and references, new methods using more efficient data structures and algorithms such as BWA [23] have been applied. For IBD detection, methods that are based on enumerating all pairwise comparisons of the individuals in a population are not scalable for large populations. Among existing methods, GERMLINE is the only one that prevents all pair comparison. However, it does not necessarily grow linearly and it depends on the total number of seed matches which may grow with the relatedness among the individuals within the population. The time complexity of GERMLINE mainly depends on the expected number of matches.

**Table 1.** Comparison of different tools for searching for identical segments

| **Problem** | **Pairwise comparison** | **Seed and extension** | **Global index** |
| --- | --- | --- | --- |
| Sequence comparison | Smith-Waterman | BLAST | BLAT/BWA |
| IBD search | PLINK/IBDseq | GERMLINE | RaPID |

Rather than the bottom-up seed-and-extension approach, building a top-down global index containing all pairwise matches larger than a given length seems more attractive. Such an index was not available until Richard Durbin invented the Positional Burrows-Wheeler transform (PBWT) [24]. Like BWT [25], PBWT facilitates a very fast and efficient exact matching approach among segments at same positions across the entire sample. Because of this trick, PBWT does not require pairwise comparison of all individuals and scales up linearly with the sample size. However, the major drawback of the PBWT algorithm is that it cannot tolerate genotyping or phasing errors. Long continuous matches (IBD segments) may have been interrupted by genotyping or phasing error or in rare cases by single mutations. As a result, direct application of PBWT will fail to detect those IBD segments.

In this work, we present a novel approach to find candidate IBD segments based on PBWT. We designed a randomized algorithm that first produces multiple low-resolution PBWTs on random subsets of markers, and then combines the results efficiently using interval tree data structure. Multiple PBWT runs are needed because a single run of PBWT on randomly selected markers usually will have low power and accuracy. A statistical framework is developed to guide the choice of hyperparameters of PBWT. We will present the algorithm in details, followed by simulations to benchmark RaPID against existing methods. Finally, we present an analysis of the 1000 Genomes Project data and show that it can reveal relatedness inside populations and also may provide insight into population history.

## 2. Methods

### 2.1 PBWT

PBWT [24] provides a fast method for finding all matches with a length greater than *l*, where *l* is the length of the match in terms of the number of variant sites. Given a panel of *M* sequences with *N* variant sites, it can compute all matches with the minimum length of *l* in *O*(*max*(*M N*, *number of matches*)) time. The PBWT algorithm can also find all maximal matches in *O*(*MN*). A sequence *s* has a maximal match in [*k*_1_,*k*_2_] to *x*_*i*_(*i*-th sequence in the panel)if the match *x*_*i*_[*k*_1_,*k*_2_] == *s*[*k*_1_,*k*_2_] cannot be extended and there is no longer match in the panel that includes [*k*_1_,*k*_2_]. The basic idea of PBWT is to sort the sequences by their reversed prefix at each position. The algorithm sweeps through the list of variant sites and keeps the starting positions of the matches between neighboring prefixes. The algorithm can be used to find IBD segments in large cohorts very efficiently in terms of time and space. However, as pointed out by Durbin, PBWT searches for exact matches and it does not account for genotyping or phasing errors. One may use shorter matches as seed and extend the matches, similar to the approach adopted by GERMLINE. But choosing an appropriate length as seed is not trivial. In particular, short seeds will result in many matches and the running time will increase dramatically.

### 2.2. Random Projection PBWT

In order to tolerate genotyping errors, instead of building PBWT on the original set of markers, we construct a scale-down lower-resolution PBWT using a hash function *y* = *f*(*x*) that translates a long bit-string vector *x* = (*x*_1_, *x*_2_,…, *x*_*p*_) into a single bit. Specifically, we divide the allele sites into windows of the same length, either in terms of number of allele sites or genetic distance (e.g., cM). The hash function *f* computes a representative value for a set of SNPs. The simplest hash function is just a bit at a randomly chosen position in a window. We could also use 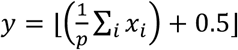. To make it more informative, we may put more emphasis onto rare variants, e.g. *y* = *sign*(*a* · (*w*⨀*x*)), where *a* and *w* are vectors of same size as *x*, *a*_*i*_’s are random sign variables (either -1 or +1), *w*_*i*_’s are nonnegative marker weights, · is the dot product, and ⨀ the element-wise product. This hash function is essentially a random projection. In this work, we chose the random bit approach for its simplicity. Figure 1 shows a schematic description of a simple example with the window size of the length 10. Note that this downsampling projection allows tolerance of errors, but may decrease the specificity of identifying true IBDs.

**Figure 1.**
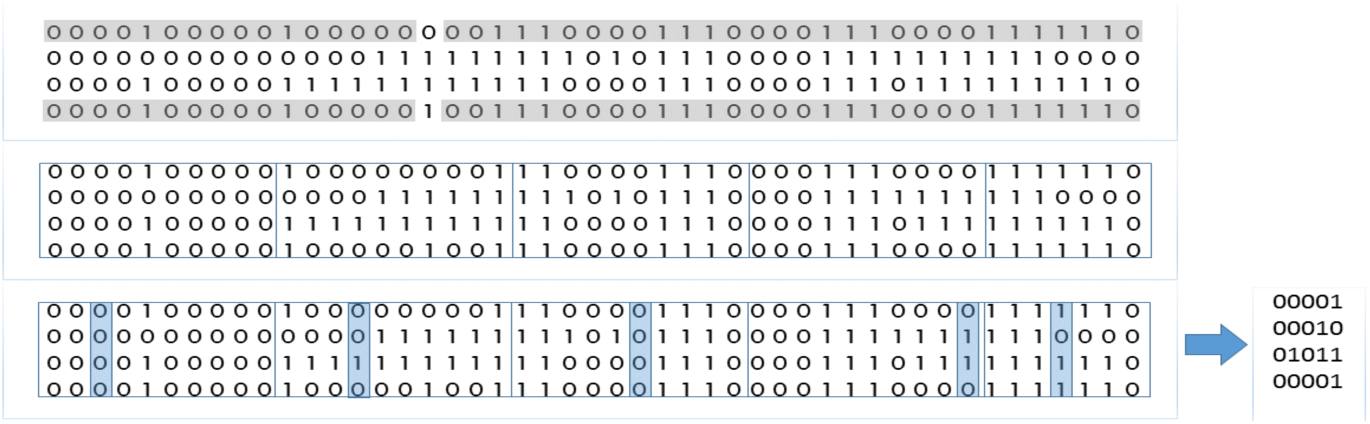
A simple example of using window size of the length 10. An allele site is selected at random for each window.

To make the approach more specific, we propose to run the random projection PBWT algorithm multiple times, each time with a different set of randomly selected variant sites. The intuition is that, because genotyping error rate is presumably low (<0.01), the variant sites with genotyping errors will only be chosen with a low probability, and thus a true IBD segment pair will have high probability to be identified in multiple runs. On the other hand, one non-IBD segment pair may be selected in some runs by pure chance, but they will not have high probability to be selected multiple times.

Specifically, we can model these probabilities as binomial distributions with relevant parameters. The parameters of RaPID are: (*r*, *w*, *c*), where *r* denotes the number of runs, *w* the window size and *c*(≤ *r*) is the minimum number of times that a match should be found in order to be considered. We run random projection *r* times, with each window containing *w* SNPs, and then run PBWT and consider any returned match with > *l* SNPs as a “hit”. The number of hits follows a binomial distribution for both true IBDs and non-IBDs. For a true IBD segment pair with identical original sequence, we assume error rate of *ε*(≪ 1), including both mutation and genotyping error rate, the binomial probability in each run is 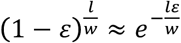, or *X_T_*~binomial (r, *e*^−*lε*/*w*^). For a random pair of segments, the probability of having a *l*/*w*-window hit is 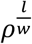, where *ρ* is the probability that a randomly chosen pair of chromosomes would share the projected sequencing in a window, or *X*_*F*_∼binomial 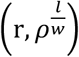. The success of random projection PBWT relies on the choice of a parameter c, such that the power Pr(*X*_*T*_ ≥ *c*) is high, while the expected number of false positives 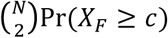 is low. Note that *ρ* is a population genetics parameter determined by haplotype frequency, which varies from region to region. *ε* is determined mainly by genotype error rate as mutation rates is typically much lower. *ε* is in the range of 0.001-0.01

An efficient strategy to choose the parameters is to estimate the number of runs and minimum number of passes so that a large range of subsampling size will result in high true positive and low false positive probabilities. It is often reasonable to set 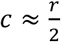, where binomial probability density distribution is far away from the boundary cases (*c* = 0 or *c* = 1). The value of *r* should be large enough to show clear separation between true positives and false positvies, and at the same time small enough to limit the running time. Figure 2a shows the true/false positives with growing for a population with a random haplotype matching probability of 0.99%.

**Figure 2.**
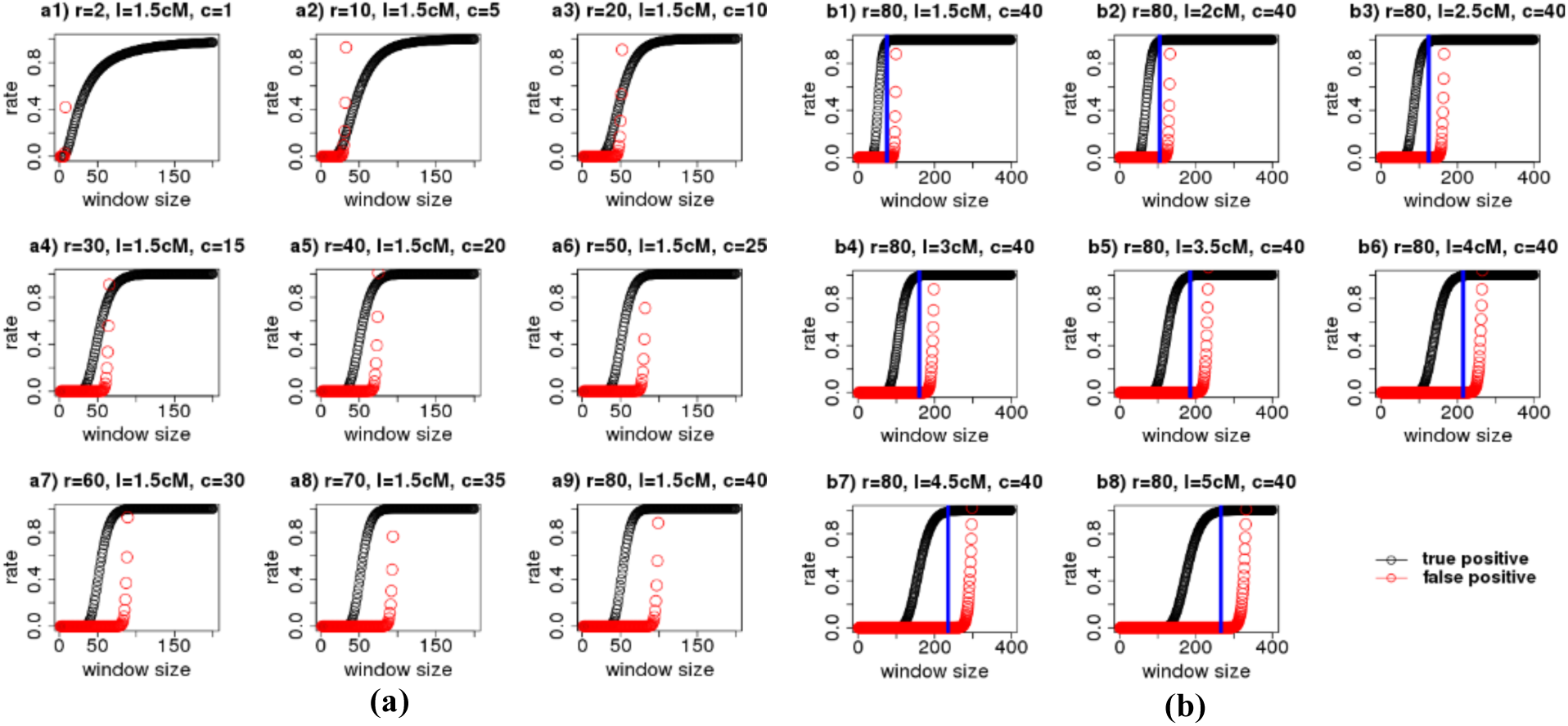
(a) Expected true/false positive rates of RaPID based on the binomial cumulative distribution function for detecting IBD segments with growing *r* among 4000 haplotypes with genotype error rate of 0.0025 and probability of random match of haplotypes of 0.99. (b) Expected true/false positive rates of RaPID based on the binomial cumulative distribution function (r = 80) for detecting IBD segments with different lengths among 4000 haplotypes. Blue vertical lines show the selected window size in our simulations.

Figure 2b (right) shows the true/false positive probabilities based on the binomial distribution of finding true and non-IBDs. As shown in the figure, the window size should set larger as we search for larger IBDs if r and c remains constant. For 1.5 cM, the optimal window size would be between 50 and 90, while for 5 cM the window size should be set between 200 and 300.

Each run of PBWT algorithm will produce a list of the matches that exceed a given length. The length is defined in terms of consecutive variant sites. Assuming the variant sites are distributed equally across the chromosomes, then the average number of variant sites to gain the minimum length can be computed easily by 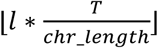, where *l* is the minimum IBD length in Mbps or cM and *T* denotes the total number of variant sites in the chromosome. chr_length denotes the length of the chromosome in Mbps or cM. If the variant sites are not distributed equally, then the reported IBD segments will not necessarily correspond to the desired minimum length and some segments will be missed. In order to prevent missing IBD segments, the actual minimum number of variant sites to cover the minimum length given in cM or Mbps can be computed. For the first allele site, the minimum number of consecutive allele sites is computed to gain the minimum length in cM or Mbps. The minimum number and the covered length can then be used for the next allele site. The distance of the first and the second sites is subtracted from the covered length and more sites from the right side are added to get the minimum length. We loop over all allele sites and the minimum number of SNPs starting from each site is computed. The minimum value is then considered as the minimum length in terms of SNPs. Finally, the variant sites at the beginning and end of reported IBD segments can be mapped into their corresponding genomic locations to filter the IBD segments shorter than the desired length.

### 2.3 Merging of PBWT Runs

In order to prevent false positives, only those hits are considered as true that occur at least *c* times. However, it is very unlikely that the starting and ending positions of two different hits from different runs are exactly the same. Instead of checking the exact starting and ending positions, we consider the overlap between two hits or intervals. The outputs of different runs should also be merged together even if the value *c* is set to 1 to remove redundant hits from different runs.

Each output of the PBWT run contains the indices of two haplotypes *k*_1_, *k*_2_, starting and ending positions of the match. To filter and merge the intervals efficiently, interval trees are used. An interval tree can be constructed in *O*(*t log t*) for *t* intervals and queried in *O*(*h* + *log t*), where *h* is the number of overlapping intervals. The large number of exact matches during each run requires an efficient method to merge the results. In order to compare the outputs, we sort the hits from each run by their indices using *Counting Sort.* It has time and space complexity of *O*(*n* + *k*), where *n* is the number of entries and *k* the maximum key value*. Counting sort* is highly time- and space-efficient for sorting hits. Note that the maximum key value is the total number of individuals or haplotypes while the number of hits is usually larger than the total number of haplotypes.

While merging the outputs of multiple runs, memory usage is crucial, since each output may contain millions of entries. We used pointers to extract the hits simultaneously from different runs that are stored in separate files. Assume the outputs are sorted based on *k*_1_ and *k*_2_ and *k*_1_ < *k*_2_. For each PBWT output *i*, a pointer variable *p*_*i*_ is used to point to the current hit that is being processed. A global variable *m* contains the minimum value of (*k*_1_, *K*_2_) pair. *R*_*i*_ denotes the results of the *i*-th output of PBWT. *R*_*i*_[*p*_*i*_] is added to the current interval tree and a set *S*, as long as *R*_*i*_[*p*_*i*_] is equal to and: is increased by 1. If none of the *p*_*i*_ variables changes then each element in *S* is searched in the interval tree. A hit is stored if the number of overlapped intervals exceeds the given threshold *c*. The remaining hits are then discarded and the variable *m* is updated. Figure 3 illustrates the pseudo-code for merging the outputs of multiple PBWT runs. The procedure *mergePBWTs* gets a set of PBWT outputs sorted by their indices and the parameter *c*. It computes the hits that occur at least *c* times out of *r* different PBWT runs.

**Figure 3.**
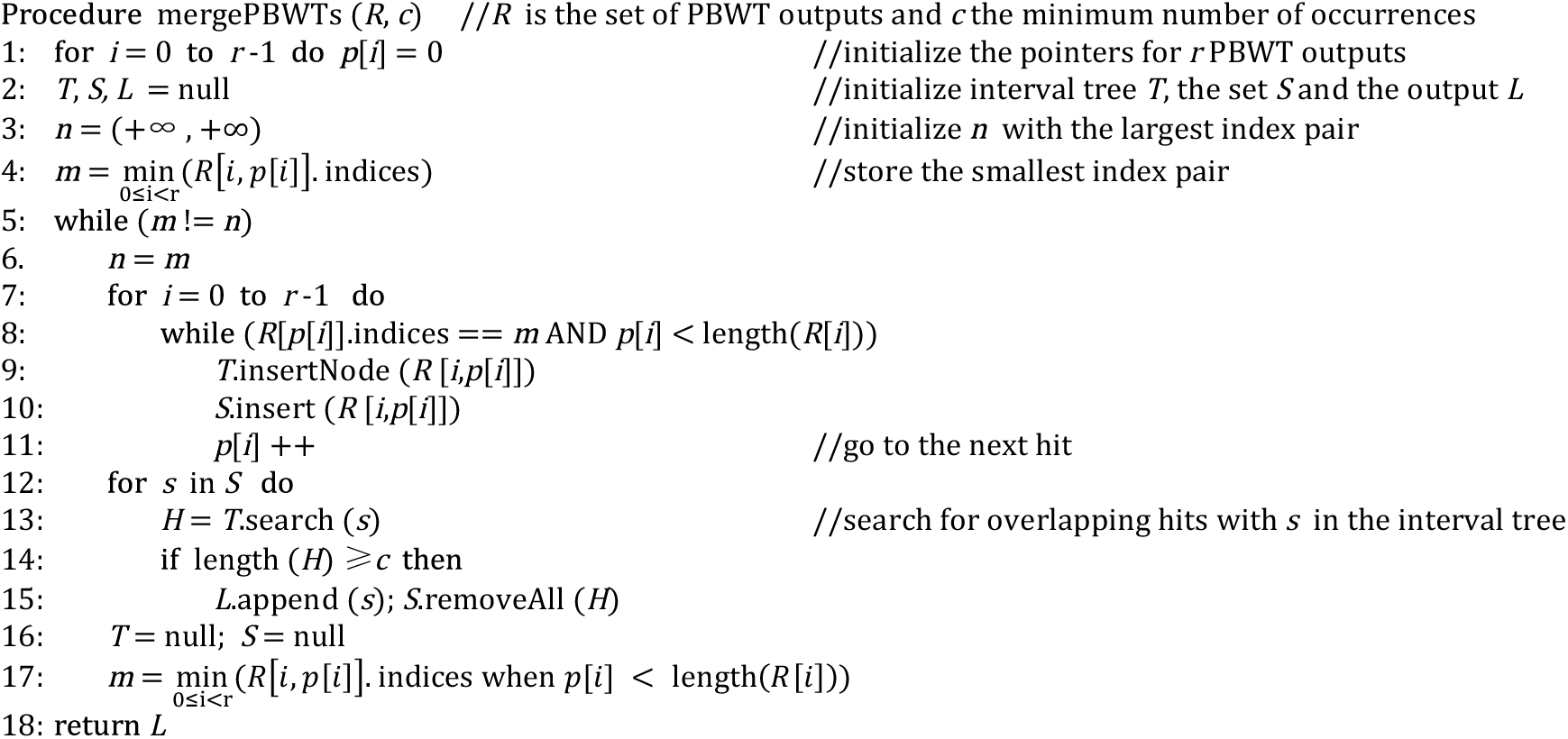
Pseudo-code for merging the results of multiple PBWT outputs.

## 3. Results

To benchmark the detection power, accuracy, and efficiency of our tool, we compared it with GERMLINE and IBDseq. PARENTE2 was not included in our bench-marking mainly due to the required large memory. We were not able to run PARENTE2 on our simulated data of 2000 individuals on a PC with 128GB memory. We ran PARENTE2 on 10% of our simulated population and it used ~10 GB of memory. The running time was also significantly worse than GERMLINE and our method. It took 5,480 seconds for 10% of the simulated data. PARENTE2 scales linearly with the chromosome length, but scales quadratically with the population size. As a results we may expect approximately 548,000 seconds or 152 hours to run the program on our entire simulated data. The major advantage of PARENTE2 over GERMLINE and our approach is that it does not require phased genotypes. We also applied RaPID on a large simulated population with 50k individuals. Finally, we applied RaPID on real data from the 1000 Genome project data.

### 3.1 Simulation

To compare with existing tools, we generated 4000 haplotypes of the length 10 Mbps and their ancestry trees using macs simulator [26], assuming a population with a history similar to that of the current Europeans with a mutation rate of 1.3x 10^−8^ and a constant recombination rate (1 cM per 1 Mbps) [19]. To simulate the genotyping error in the generated haplotypes, we inserted an error rate of 0.0025 for each haplotype as in [19], corresponding to genotyping quality of 26 in standard VCF files. The true IBDs were determined as [19]: we sampled the ancestry trees at every 5 kbps and if the most recent common ancestor of a pair of individuals remains constant, the corresponding segment of the pair was considered as true IBD. These were achieved by using the Dendropy python library. To validate the capability of our tool in handling very large sample sizes, we also generated 100k haplotypes of length 10 Mbps with a mutation and recombination rate of 0.001 to benchmark the performance of RaPID for very large cohorts.

### 3.2 Benchmarking

In order to evaluate the correctness of detected IBDs, we computed accuracy and power of RaPID, GERMLINE and IBDSeq for simulated data of 2000 individuals. Accuracy is defined as the percentage of correctly detected IBDs with at least 50% overlap among the total number of reported IBDs. Power is defined as the average proporotion of correctly detected IBD segments. Subsequently, we demonstrate the efficiency of RaPID regarding the running time with increasing number of haplotypes.

### 3.2.1 Accuracy and Power

The parameters of RaPID were set as described in the method section. We set the number of runs *r*=80 and the minimum number of passes *c*=40. The window size was then estimated based on the probability distributions of true/false positives. The window sizes were set to 75, 105, 125, 160, 185, 215, 235 and 265 to detect IBDs with a minimum length of 1.5 to 5 cM, respectively. Accuracy and power of RaPID, GERMLINE and IBDseq are shown in Figure 4. RaPID can achieve higher accuracy than GERMLINE and comparable results to IBDseq for shorter IBDs. GERMLINE is able to find most of the IBDs, however for shorter IBDs the results are not accurate. The accuracy of all methods increases by increasing the minimum length of IBD. Since the probability of very long identical segments by chance decreases, we also expect higher accuracy value for longer IBD segments.

**Figure 4.**
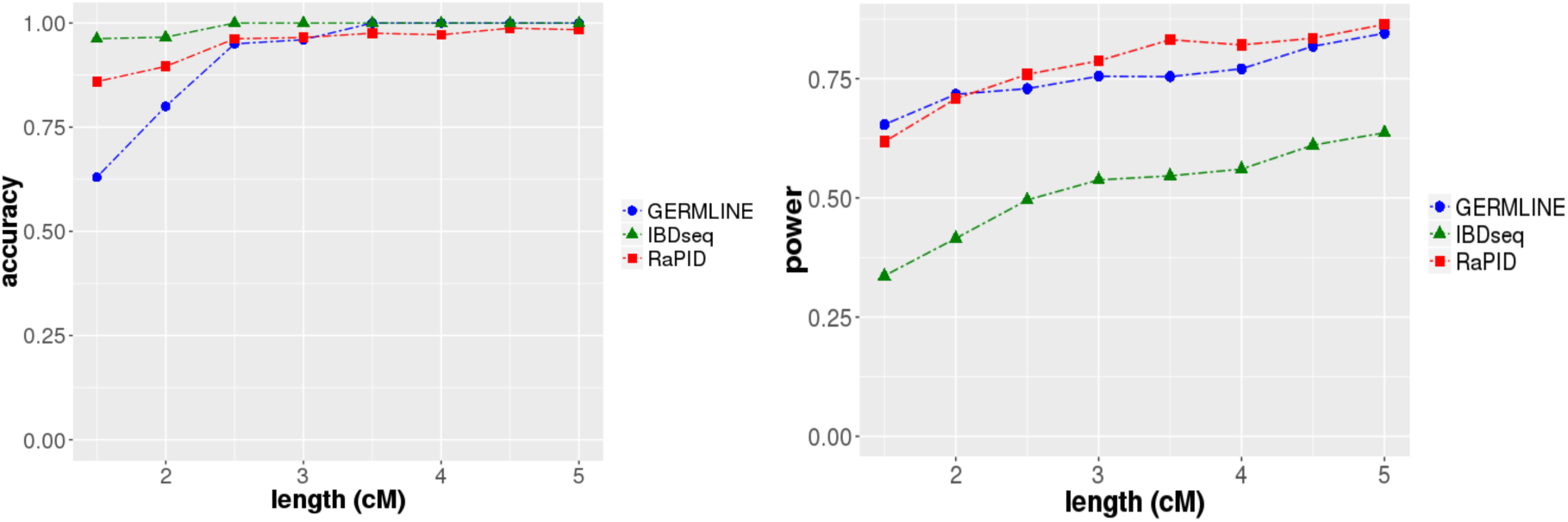
Accuracy and Power of GERMLINE, IBDseq and RaPID with the increasing length of IBD among 2000 individuals. Results are binned by segment size: bins are extended 0.2 cM on the right side for 1.5 and 2 cM; and 1 cM for x axis values > 2.

### 3.2.2 Running Time

IBD detection tools that require pairwise comparison of all haplotypes are generally not scalable for large number of haplotypes. Both IBDseq and PARENTE2 perform pairwise comparison for genotype data. As a results, they are very slow compared to GERMLINE and RaPID. Table 2 shows the running time of different methods for 2000 individual with a chromosome length of 10 Mbps. In our experiments, the running time for RaPID was more than 100 times less than that of GERMLINE for detecting IBDs with a minimum length of 3 cM and almost 25 times less for IBDs with a minimum length of 1.5 cM. The running time of RaPID can be impacted by the selected parameters. The minimum length of IBDs and the level of heterogeneity in the sample are the most important factors in determining the running time of RaPID.

**Table 2.** Running time for finding IBDs in the simulated data containing 2000 individuals using different methods

| Method | Size (cM) | Accuracy | Power | Running Time |
| --- | --- | --- | --- | --- |
| RaPID | 1.5 | 0.86 | 0.62 | 23 seconds |
|  | 3 | 0.97 | 0.79 | 5 seconds |
| GERMLINE | 1.5 | 0.63 | 0.65 | 8.72 minutes |
|  | 3 | 0.96 | 0.75 |  |
| IBDseq | 1.5 | 0.96 | 0.34 | 12.08 hours |
|  | 3 | 1.0 | 0.54 |  |
| PARENTE2 | - | - | - | 6 days* |

\*The running time for PARENTE2 is estimated based on a run on 10% of the simulated data.

We also compared how the running times of RaPID, GERMLINE and IBDseq grow with increasing population size to find IBDs with a minimum length of 1.5 cM. To compare the growth of running time, the tools were run on increasingly larger sets of haplotypes up to 4000. We set the minimum length for GERMLINE to 1.5 cM (-min_m =1.5). As depicted in Figure 5, RaPID grows linearly with the increasing number of samples, while GERMLINE shows a super linear behavior.

**Figure 5.**
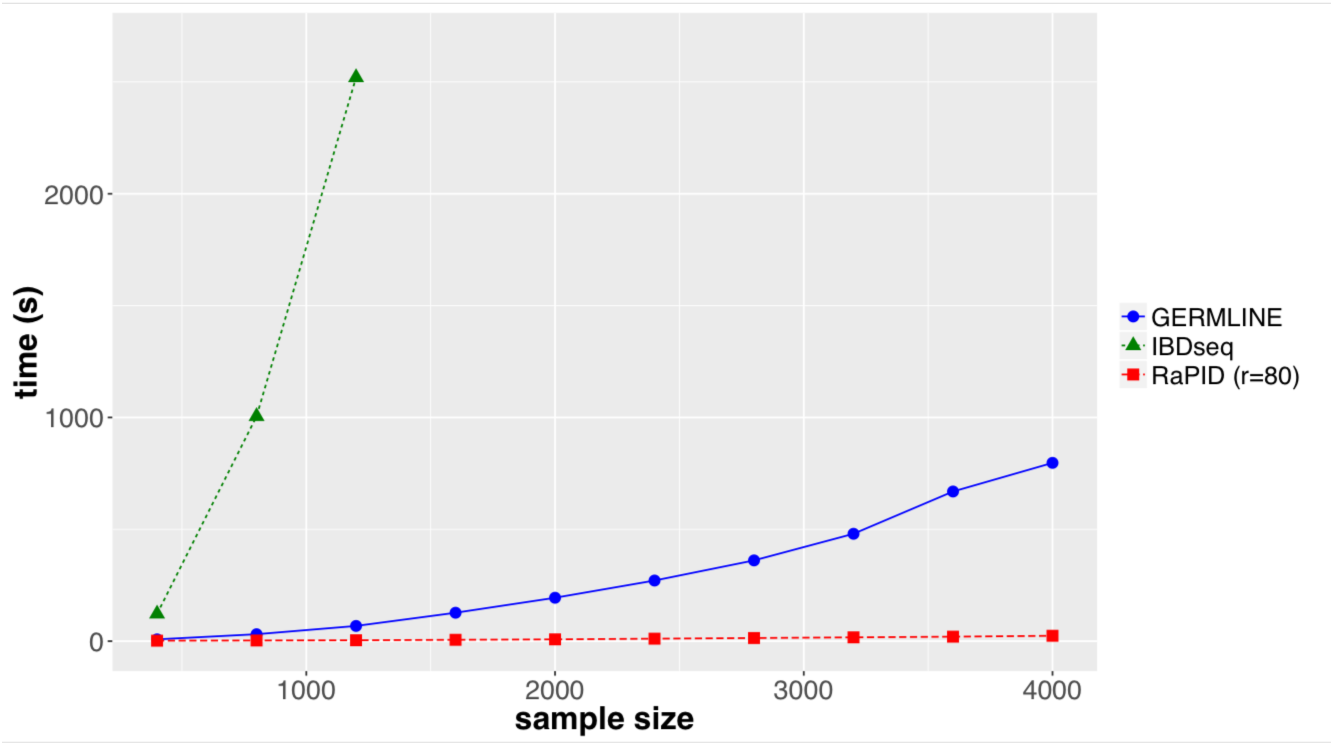
Comparison of running time of GERMLINE, IBDseq and RaPID with increasing number of haplotypes for IBDs with a minimum length of 1.5 cM.

### 3.3 Applying RaPID on 100,000 haplotypes

In order to verify the ability of RaPID to detect IBDs in a large cohort, we applied it on 100k simulated haplotypes. RaPID was applied to detect the IBDs with the minimum length of 1.5 Mbps among all of the haplotypes. The program returned the results within 10 minutes. Since extracting all of the true IBDs based on the ancestry tree generated by the simulator was not feasible in a reasonable time, we compared only the true IBDs that were shared between the first 100 haplotypes and the whole population. The power of RaPID was 100% and the accuracy 65% for IBDs with a minimum length of 1.5 cM. RaPID is the only tool that can handle hundreds of thousands of individuals in a reasonable time on a single CPU. Based on our experiments, we estimate that RaPID can handle 1 million haplotypes of the same length within few hours.

### 3.4 Applying RaPID on the 1000 Genome Project Data

To demonstrate the utility of RaPID for real data, we applied it on the 1000 Genome Project data [1]. The 1000 Genome Project provides the largest publicly available catalogue of human variation and genotype data. The phase 3 data set includes 2504 individuals from 26 populations. The execution time for RaPID was around 2 hours for detecting IBDs with a minimum length of 1.5 cM and 1 hour for IBDs with a minimum length of 5 cM for all 22 autosomal chromosomes. We downloaded the phased genotype data (in VCF format) of the 1000 Genome project^†^. We then extracted the bi-allelic sites and applied RaPID on the phased data of the autosomal chromosomes. We computed the kinship based on the length of shared IBD segments. The kinship coefficients were computed by summing the lengths of the autosomal IBD segments and dividing by four times the length of the autosomal chromosomes, as did in ftp://ftp-trace.ncbi.nih.gov/1000genomes/ftp/release/20130502/supporting/ibd_by_pair/20150129_IBD_segment_methods.pdf. Figure 6 shows the computed kinship among the populations based on the detected IBDs with a minimum length of 1.5 and 5 cM in logarithmic scale. Recently, Fedorova et al [27] have analyzed the phase 1 data using rare variant clusters (RVC) to study the genetic relationships among different populations. The detected relationships among different populations using RaPID were congruent to Fedorova et al’s discovery.

**Figure 6.**
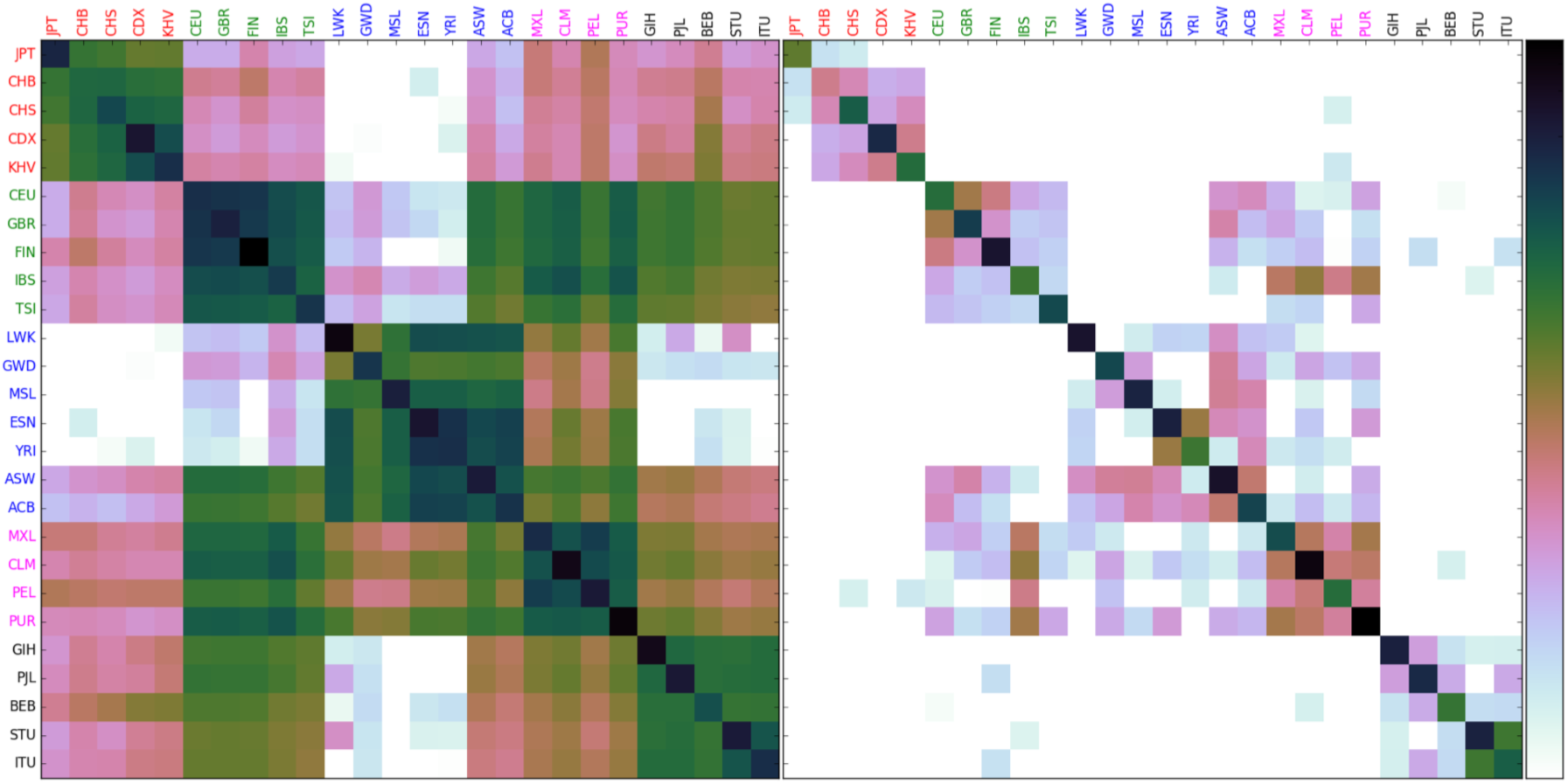
Kinship (in log scale) among 26 populations from the 1000 Genome Project based on the detected IBDs with a minimum length of 1.5 (left) and 5 cM (right). Maximum kinship value for 5 cM is 0.0017 and for 1.5 cM is 0.0037 **East Asian (in red)**: Han Chinese in Beijing (CHB), Japanese in Tokyo (JPT), Southern Han Chinese (CHS), Chinese Dai in Xishuangbanna (CDX), Kinh in Ho Chi Minh City (KHV) **European (in green)**: Utah Residents (CEU), Toscani in Italy (TSI), Finnish (FIN), British in England and Scotland (GBR), Iberian Population in Spain (IBS) **African (in blue)**: Yoruba in Ibadan (YRI), Luhya in Webuye Kenya (LWK), Gambian (GWD), Mende in Sierra Leon (MSL), Esan in Nigeria (ESN), Americans of African Ancestry in SW USA (ASW), African Caribbeans in Barbados (ACB) **Ad Mixed American (in magenta)**: Mexican from LA (MXL), Puerto Ricans (PUR), Colombians from Medellin (CLM), Peruvians from Lima (PEL) **South Asian (in black)**: Gujarati Indian from Houston (GIH), Punjabi from Lahore (PJL), Bengali from Bangladesh (BEB), Sri Lankan Tamil from the UK (STU), Indian Telugu from the UK (ITU).

As expected, within-population kinship is always higher than the between-population kinship. In Asia, Chinese Dai in Xishuangbanna (CDX) and Japanese people (JPT) have the highest within-population kinship. In Europe, the highest kinship value is observed among Finnish people. In Africa, LWK (Luhya in Webuye, Kenya) has the highest within-population kinship value. In America, Puerto Ricans have the highest kinship. In South Asia, Gujarati Indians (GIH) have the highest kinship. ITU (Indian Telgu from UK) and STU (Sir Lanken Tamil from UK) have the highest kinship values among South Asian populations.

While most of the detected IBDs with a minimum length of 5 cM are within the same population, there are also some shared IBD segments between the populations from the same continent. However, some populations do not share any or only very few IBD segments within their corresponding continents. At 5 cM level, Japanese people are almost separated from other populations in Asia. The lowest kinship value among Europeans is between Finnish and Italian populations (TSI). According to [27] the lowest number of shared RVC is also observed among Finish and South European populations. The highest kinship among Europeans is between people from Utah (CEU) and Britain (GBR). The highest number of RVC among European populations was also reported to be between CEU and GBR [27]. The highest kinship among African population is between ESN (Easn in Nigera) and YRI (Yoruba in Ibdadan, Nigera). Furthermore, the kinship values at 5 cM reveal that American populations are related to the Iberian population in Spain (IBS).

Shorter IBD segments reveal more ancient relationships among different populations. Inter-continental kinship values based on the computed IBDs with a minimum length of 1.5 cM are also low, but they may still reveal interesting ancient relatedness. Finnish people have the highest kinship values with East Asian people compared to the other European populations. The kinship values among Asian and African people are close to zero. American populations share IBD segments with Asian populations. Among American populations, Peruvians (PEL) are the most related population to Asian people followed by Mexicans (MXL). Bangali people from Bangladesh (BEB) are the most related population to East Asian people, especially South Eastern people, compared to the other populations from South Asia.

## 4. Discussions

In this paper, we present a fast method to detect IBD segments of a given length in very large cohorts and investigate the relatedness in large panels which may not be feasible using current methods. The proposed method RaPID searches for consecutive haplotype matches allowing genotyping error. Based on our experiments, it can detect IBDs with a minimum length of more than 1 cM very efficiently while maintaining comparable accuracy and power to GERMLINE and IBDseq. RaPID can handle hundreds of thousands or millions of individuals in a reasonable time by taking advantage of the underlying population structure and targeting the IBD segments of the given length. Therefore, RaPID is currently the only IBD detection method for large biobank-scale population genotype data.

In our analysis of the phased data of the 1000 Genome project, the detected IBDs with a minimum length of 5 cM were almost from the same population. More IBD segments with a minimum length of 1.5 cM were detected including IBDs within the same super population (or continent). A few inter-continental IBD segments were also detected at 1.5 cM level. By tuning the target IBD segment length, RaPID can reveal population history events in different time scale.

While we presented the proof-of-concept version of RaPID, there are potential improvements for future works. We will extend RaPID for unphased genotype data. Current RaPID is based on phased data, since it is based on the PBWT index for haplotypes. We can first phase genotype data by computational methods. Because the high efficiency of RaPID, we can run RaPID over multiple alternative phasing results of the same genotype data so that our method is tolerant to phasing errors.

Another potential extension is to improve the detection of very short IBD segments. Identifying IBD with length below 1 cM is a difficult problem because the distinction between true positives and false positives can be blurred. Our results revealed that short IBD segments are actually very informative for ancient population events. Because RaPID can be optimized to identify IBD segments of a certain length range, our tools can be more sensitive in identifying shorter IBDs.

## Availability

The most recent version of RaPID is freely available at https://github.com/ZhiGroup/RaPID. Detailed information of parameters used in the benchmarking and the application to the 1000 Genome Project Data and the corresponding version are available at http://genome.ucf.edu/RaPID.

## Acknowledgment

We would like to thank Dr. Sharon Browning from the Department of Biostatistics at the University of Washington for her assistance and valuable comments on the benchmarking data.

## Footnotes

† ftp://ftp-trace.ncbi.nih.gov/1000genomes/ftp/release/20130502/

